# Manganese homeostasis modulates glucan and chitin unmasking in the opportunistic yeast *Candida albicans*

**DOI:** 10.1101/2025.03.24.644998

**Authors:** Manon Henry, Maria Khouas, Gabriel Théberge-Julien, Antony T. Vincent, Louis Villeneuve, Éric Rhéaume, Jean-Claude Tardif, Adnane Sellam

**Affiliations:** Montreal Heart Institute, Université de Montréal, Montréal, QC, Canada; Department of Animal Sciences, Université Laval, Quebec City, QC, Canada; Department of Medicine, Université de Montréal, Montréal, QC, Canada; Department of Microbiology, Infectious Diseases and Immunology, Faculty of Medicine, Université de Montréal, Montréal, QC, Canada

**Keywords:** *Candida albicans*, Manganese homeostasis, Smf12 transporter, Cell wall unmasking

## Abstract

*Candida albicans* is a commensal fungus and also the most prevalent human fungal pathogen. The ability of this opportunistic yeast to acquire and maintain homeostatic levels of manganese (Mn), particularly in the metal-limited host environment, is an important determinant of its fitness. Recent studies have underscored the importance of Mn acquisition through members of Smf transporters, in *C. albicans* virulence and its ability to withstand various stresses. In the present study, we undertook transcriptional profiling in the mutant of the Mn transporter Smf12 under restricted Mn availability to identify processes that are directly affected in defective Mn uptake. Our analysis revealed that *smf12* displayed a transcriptional pattern suggestive of a cell wall defect, with many transcripts associated with cell wall biogenesis being differentially regulated. *smf12* together with *smf11*, a mutant of the closest homolog of Smf12, exhibited hypersensitivity to cell wall stressors and an altered cell wall ultrastructure. The *smf* mutants also exhibited unmasking of both β-glucan and chitin, which unexpectedly resulted in a decreased rate of phagocytosis by macrophages, suggesting impaired recognition or internalization—an observation that challenges the prevailing paradigm. Furthermore, we showed that Mn-mediated unmasking of β-glucan required modulation of glucanase activity and was not mediated through the calcineurin pathway. This study uncovers a novel role for Mn in maintaining cell wall integrity and modulating the exposure of fungal antigenic determinants, further emphasizing the critical role of this metal in supporting the opportunistic nature of *C. albicans*.

## Introduction

The availability of transition metals, such as iron (Fe), copper (Cu), and zinc (Zn), is tightly regulated within the human host. To limit the proliferation of microbial pathogens, the host employs a strategy known as nutritional immunity, which involves sequestering these essential metals or directing their toxicity against pathogens ^1^. However, many pathogens have evolved diverse strategies to bypass nutritional immunity, allowing them to survive and infect the host. Fungal pathogens, in particular, have developed sophisticated regulatory networks to modulate both metal acquisition and detoxification, thereby maintaining homeostasis of these essential micronutrients. For instance, in *Candida albicans* and other human fungal pathogens, Fe acquisition is facilitated by the high-affinity reductive and siderophore transport systems, as well as hemoglobin-Fe utilization, all of which are essential for virulence ^2–5^. Similarly, Cu and Zn are internalized by specific transporters regulated at the transcriptional levels by the transcription factors Mac1 and Csr1, respectively, which are also essential for fungal fitness *in vivo* ^6–8^. Although significant attention has been given to the mechanisms by which fungi acquire Fe, Cu, and Zn, the role of Mn in infectious processes and the cellular mechanisms underlying Mn homeostasis in fungal cells remain partially understood.

*C. albicans* is a commensal fungus and an important member of the human intestinal and vaginal microbiota. Under certain conditions, including immune suppression and dysbiosis, this opportunistic yeast can cause serious infections, such as fungal sepsis, which is associated with high mortality rates ^9^. While significant attention has been given to the mechanisms of Fe and Cu acquisition and their role in fungal fitness, less focus has been devoted to manganese (Mn). Recently, two independent studies have highlighted the role of manganese (Mn) homeostasis in the biology and the fitness of this prevalent human fungal pathogen ^10,11^. Mn uptake via the NRAMP family transporters Smf12 and Smf13 has been shown to be essential for the development of invasive hyphae and to fully support the activity of Mn-dependent superoxide dismutases—critical enzymes that protect *C. albicans* cells from oxidative stress ^10–12^. Furthermore, Mn has been shown to modulate antifungal sensitivity by regulating antifungal efflux and ergosterol homeostasis ^11^. Given the critical role of Mn in these virulence-related processes, both *smf12* and *smf13* mutants displayed an altered virulence in various infection models ^10,11^.

Mn also serves as a cofactor for several metalloenzymes, including mannosyltransferases, which are essential for mannan biosynthesis and the generation of cell wall mannoproteins ^13,14^. Inactivation of Mn transporters, such as Smf12 or the P-type ATPase Mn^2+^/Ca^2+^transporter Pmr1, results in a shortening of glycosyl residues linked to proteins ^11,13^. Since mannoproteins are immunogenic epitopes that decorate the outer fungal cell wall, Mn homeostasis likely plays a pivotal role in modulating immune recognition. Indeed, under Mn deprivation, *C. albicans* cells unmask β-glucan, a major polysaccharide of the inner cell wall layer and another important fungal antigenic determinant ^15^. This unmasking is likely due to the reduced density of mannan fibrils in the outer layer of the cell wall ^13^. In contrast, Fe starvation leads to the opposite effect, where β-glucan is masked, reducing consequently the recognition by phagocytic macrophages ^15^. This Fe-mediated masking is regulated by the cAMP-Protein Kinase A signaling pathway, along with the Fe transceptor Ftr1 and the Fe-responsive transcription factor Sef1 ^15^. In contrast to Fe, however, the mechanism by which β-glucan is unmasked under Mn limitation and whether this enhances immune cell recognition remains unexplored.

In the current study, we conducted transcriptional profiling of the *smf12* mutant in *C. albicans* under restricted Mn availability to identify processes that are directly affected by a defect in Mn internalization. *smf12* mutant exhibited a transcriptional pattern reminiscent of a cell wall defect, with many transcripts related to cell wall biogenesis being differentially regulated. Accordingly, *smf12* together with *smf11*, a mutant of the closest homolog of Smf12, and their corresponding double mutant (*smf12 smf11*), exhibited hypersensitivity to cell wall stressors and altered cell wall ultrastructure. Consistent with the observation that Mn depletion led to increased exposure of β-glucan ^15^, the *smf12* mutant also exhibited this effect, along with unmasking of the basal chitin layer of the inner cell wall. Intriguingly, unmasking of either β-glucan or chitin in *smf* mutants led to a decreased rate of phagocytosis suggesting a reduced recognition or internalization by macrophages, which challenges previous paradigm. Moreover, we showed that Mn-mediated unmasking of β-glucan required the modulation of the glucanase activity and was not signaled through the calcineurin pathway.

## Materials and methods

### Strains and growth conditions

The fungal strains and the PCR primers used in this study are listed in **Supplementary Table S1**. *C. albicans* clinical strain SC5314 ^16^ and its derivatives were routinely maintained at 30°C on synthetic complete (SC; 1.7% yeast nitrogen base, 0.5% ammonium sulfate, 2% dextrose, 0.2% amino acid, with 50 µg/ml uridine). *smf* deletion mutants (*smf12*, *smf11* and the double mutant *smf12 smf11*) were constructed from SN148 strain ^17^ by replacing the entire ORF with a PCR-disruption cassette amplified from the pFA plasmids ^18^. A wild-type control strain (SN148-CIp20) was created by reintroducing *URA3* and *HIS1* in SN148 using the integrative Cip20 plasmid ^19^.

### Cell wall stress assays

Overnight cultures on Mn-free synthetic complete growth medium (SC-Mn) were diluted to an OD_600_ of 0.1 and 5-fold serial dilutions were prepared using the same medium. A total of 4 μl of each dilution was spotted on either SC-Mn or SC+Mn (SC-Mn with 1 mM MnCl_2_) agar medium containing 0.2 µg/ml caspofungin or 25 µg/ml calcofluor white. Plates were incubated at 30°C for 2 days.

### Expression analysis by RNA-seq

RNA-seq was conducted as previously described by Henry *et al.* ^11^. Briefly, overnight cultures of *smf12* strain were diluted to an OD_600_ of 0.1 in 60 ml of fresh SC-Mn medium and grown at 30°C under agitation to early logarithmic phase (OD_600_=0.4). Cultures were then either supplemented (SC+Mn) or not with 1 mM MnCl_2_ and incubated at 30°C for 90 min. Cell pellets were collected by centrifugation and total RNA was extracted using an RNAeasy purification kit (Qiagen) as previously described ^20^. cDNA libraries were prepared using the NEBNext Ultra II RNA Library Prep Kit and sequencing was performed using an Illumina NovaSeq 6000 sequencing system. Gene ontology (GO) analysis was performed using GO Term Finder of the Candida Genome Database ^21^. The GSEA (Gene Set Enrichment Analysis) pre-ranked tool (http://www.broadinstitute.org/gsea/) was used to determine the statistical significance of correlations between the *C. albicans smf12* transcriptome and different omics datasets as previously described ^22^. Differentially expressed transcripts in **Supplementary Table S2** were identified using Welch’s t-test with a false-discovery rate (FDR) of 5% and 1.5-fold enrichment cut-off. All RNA-seq data are available at the GEO database (https://www.ncbi.nlm.nih.gov/geo/) with the accession number GSE283948.

### Quantification of **β**-glucan, chitin, and mannan by flow cytometry

*C. albicans* cells were grown in SC-Mn and SC+Mn until reaching the exponential phase and were centrifuged and fixed for 20 min using 4% paraformaldehyde (w/v, final concentration). Cells were washed three times in FACS buffer (PBS, 0.5mM EDTA, 0.1% BSA). A total of 100 µl cell suspension (OD = 1) was treated as follows. For β-glucan exposure, cells were stained using 5 ng Dectin-1 conjugated to IgG1 Fc domain (fc-hdec1a-2, InvivoGen) for one hour at 4°C. Cells were then washed three times in FACS buffer and incubated for one hour at 4°C with IgG Fc secondary antibody conjugated to TRITC (Invitrogen, A18822). For mannan and chitin exposures, cells were incubated for one hour at 4°C with Concanavalin A conjugated to Alexa-488 (Invitrogen, C11252) and wheat germ agglutinin (WGA) conjugated to Alexa-488 (Invitrogen, W11261), respectively. Cells were then washed three times with FACS buffer and analyzed on a BD FACSymphony A1 Cell Analyzer. Fixed cells were identified by FSC/SSC using unstained cells to adjust voltages. The mean fluorescence intensity (MFI) of stained cells was measured for the TRITC and Alexa Fluor 488 channels. Data were collected for three biological replicates for each condition and analyzed with the BD FACSDiva Software v.9.0.2. Graphs present the percentage of the MFI of 10,000 events normalized to the WT in SC+Mn.

### Phagocytosis assay and cytokines production quantification

Phagocytosis was quantified as described by Carneiro *et al.* ^23^. Briefly, yeast cells were grown in either SC-Mn or SC+Mn to reach the exponential phase and were then fixed for 20 min using 4% formaldehyde and 0.2% Triton mixture. Fixed cells were stained with SytoxGreen 1mM for one hour and co-cultured with macrophages J774A.1 MOI (5:1) in DMEM growth medium at 37°C and 5% CO_2_. To discriminate engulfed yeasts from those attached to macrophages, we take benefit of SYTOX-green quenching by propidium iodide (PI) (6 mg/ml) for 10 min. While phagocytosed yeasts are protected from quenching by macrophage membrane, non-phagocyted or attached yeast are stained with PI. After one hour incubation, non-phagocyted yeast were stained with propidium iodide (6 mg/ml) for 10 min and quantification was made by flow cytometry. J774A.1 cells were gated by FSC/SSC and fluorescence minus one controls (FMOC) were used to define the quadrants identifying phagocytic populations in a PI/SytoxGreen dot plot ^23^. A total of 10,000 events per condition were measured and three biological replicates were considered for each condition. TNF-α and IL-6 levels in the co-culture media were measured using the mouse TNF-α (Quantikine, MTA00B-1) and IL-6 (Quantikine, RM6000B-1) ELISA kits, respectively, according to the manufacturer’s instructions.

### Microscopy

Transmission electron microscopy (TEM) was used for cell wall ultrastructure imaging (Imaging-Microscopy Platform, IBIS, Université Laval, Quebec City). Yeast cells were grown in SC+Mn or SC-Mn liquid media to reach the exponential phase and were then fixed overnight at 4°C using a mixture of 2% glutaraldehyde and 1.6% paraformaldehyde. Cells were washed three times in cacodylate buffer and were immobilized with 3% agarose, cut in a stick of 0.5 by 0.5 by 5 mm, washed twice in cacodylate buffer, and fixed in 1% OsO_4_ for 90 min at room temperature. Samples were dehydrated with grading ethanol (30%, 50%, 70%, 95%, 100%, 15 min each) and washed three times for 15 min with 100% ethanol. Samples were mixed in EPON-812 and hardener (BDMA) in a silicone mold and dried for 1 day at 37°C followed by 3 days at 60°C. Sections were cut with a Reichert-Jung Ultracut E Ultramicrotome using a 35° DiATOME diamond blade and then stained for 5 min in 2% uranyl acetate. To enhance uranyl acetate contrast, we proceed with a second stain for 3 min in 3% lead citrate. Grids were imaged on a JEM-2100plus TEM. Graph in Figure 2D represents the thickness of the inner and the outer layers in a total of 20 cells per condition and 100 measurements per cell using ImageJ ^24^. Total chitin was quantified using epi-fluorescence microscopy. *C. albicans* cells were stained with calcofluor white (10 µg/ml) and images were taken using a Biotek ® Cytation 5 high-content microscope. For each condition, fluorescence intensities of 100 cells were measured using ImageJ ^24^.

### Exo-glucanase and chitinase assays

Cells were grown in SC+Mn or SC-Mn until reaching the exponential phase and washed with 50 mM sodium acetate pH 5.5. Exo-glucanase activity was assessed according to the method previously described by Gonzalez *et al*. ^25^. Cell pellets were incubated in 200 µl sodium acetate buffer containing 1% laminarin (w/v; Sigma, L9634-1G) for 4h at 37°C. Reduced glucose released after degradation of laminarin polymer by glucanase was quantified using DNS (Sigma, D0550-100G) assay as previously described ^26^. DNS mixed with reducing sugar releases colorimetric 3-amino-nitrosalicylic acid proportionally to reduced sugar concentration. Reduced sugar concentrations in samples were determined based on a glucose standard curve (0 - 4 mg/ml). Samples were mixed in a 1:20 (sample: DNS reagent (10mg/ml)) ratio and heated for 5 min at 100°C to catalyze the reaction between sugar and DNS, and measurements were made at OD_540nm_ using Cytation 5. Chitinase activity was quantified as previously described by McCreath and Gooday ^27^. Cell pellets were incubated in 200 µl sodium acetate buffer containing 1% 4-methylumbelliferyl N-acetyl-β-D-glucosaminide (4MU) (w/v; Sigma) for two hours at 37°C. Fluorescence was measured at λ_ex_360 nm and λ_em_440 nm.

### Inductively coupled plasma-mass spectrometry

The total content of cell-associated calcium was quantified using inductively coupled plasma-mass spectrometry (ICP-MS) as previously described by Henry *et al.* ^11^.

### Statistical analyses

GraphPad Prism 10 was used for statistical analyses. Data were generated from at least three independent biological replicates and then expressed as means ± standard deviation. Statistical difference between two sets of data with a non-parametric distribution was assesses using one-way ANOVA (Tukey’s multiple comparison test). The following *p*-values were considered: * *p* < 0.05; ** *p* < 0.01; *** *p* < 0.001; **** *p* < 0.0001.

## Results

### Transcriptomic analysis of *smf12* under Mn starvation reflects a cell wall defect

In our previous work, we generated a mutant of the Mn NRAMP transporter, *SMF12*, to investigate the impact of Mn scarcity on the biology of *C. albicans*. We found that inactivation of *SMF12* resulted in attenuated virulence, increased sensitivity to azoles, and activation of the unfolded protein response (UPR) ^11^. To investigate further cellular processes affected in *smf12*, we carried out RNA-seq profiling under manganese (Mn) starvation. The transcriptional profile of *smf12* growing in the Mn-free SC medium (SC-Mn) was compared to that of the WT strain growing under similar conditions. Upregulated transcripts in *smf12* were mainly enriched in processes related to the synthesis and the processing of the three core polysaccharides of the fungal cell wall namely chitin (*CHS2*, *7*, *CDA2*, *CHT4*), glucan (*KRE1*, *9*, *UTR2*, *PNG2*) and phosphomannan (*MNN1*, *15*, *41*, *42*, *43*, *45*, *47*, *BMT3*, *4*, *6*, *7*, *KTR4*, *DFG5*, *FAV3*) (**Figure 1A-B and Supplementary Table S2**). Transcripts related to cell wall signaling proteins and transcriptional control including the Cek1 MAP kinase, the histidine kinase Chk1 and the transcription factor Crz1 were also upregulated (**Figure 1B**). This transcriptional signature is reminiscent of a situation in which the cell wall is undergoing remodeling in response to a specific perturbation. Downregulated genes were mainly enriched in the acquisition of metals including iron (*SIT1*, *CFL2*, *4*, *5*, *IRO1*, *FRE7*, *30*, *FET31*, *CSA2*, *FTR1*), zinc (*ZRT101*, *PRA1*, *CSR1*, *ZRT2*) and copper (*CTR1*) (**Figure 1C**). Genes of carbohydrate metabolism and both secreted lipases and proteases were differentially modulated in *smf12* mutant suggesting a reprogramming of metabolism in response to the deletion of this Mn transporter (**Figure 1C**). When Mn was supplemented to the growth medium, the differential modulation of most of the *smf12* transcripts described above was significantly attenuated (**Figure 1B-C**). This suggests that the transcriptional profile of *smf12* is primarily a result of impaired Mn homeostasis in this mutant.

**Figure 1.**
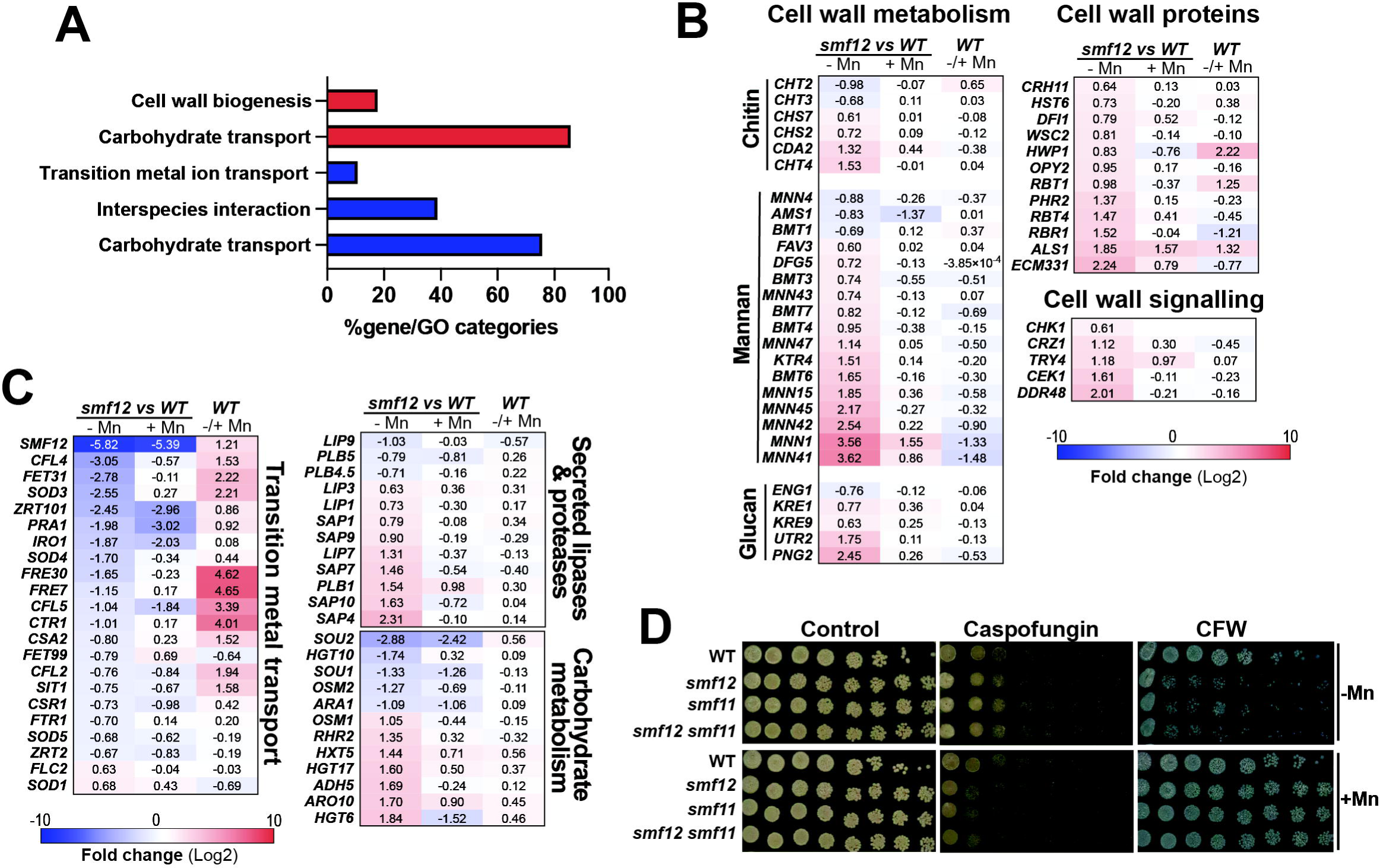
*smf12* exhibits a transcriptional signature reflecting cell wall perturbation. (**A**) GO enrichment of upregulated (red) and downregulated (blue) transcripts in *smf12* mutant growing under Mn limitation. The transcriptional profile of *smf12* growing in SC-Mn was compared to that of the WT strain growing under similar conditions. (**B-C**) Modulation of cell wall associated genes (**B**) and transcripts enriched in metals utilization and metabolism (**C**) in *smf12* mutant. Heatmaps represent differentially modulated transcript in *smf12* mutant under both Mn limitation (-Mn) and repletion (+Mn) as well as in WT in response to Mn starvation ^11^. (**D**) *smf* mutants are sensitive to cell wall perturbers in an Mn-dependant manner. Spot assay was used to assess the impact of caspofungin (0.2 µg/ml) and calcofluor white (CFW; 25 µg/ml) on the growth of WT (SN148-CIp20) and *smfs* mutants (*smf12*, *smf12* and *smf12 smf11* double mutant). Strains were serially diluted, spotted on SC-Mn and SC + Mn, and incubated for 2 days at 30°C.

To test the potential cell wall perturbation in *smf12*, we assessed the sensitivity of this mutant as well as *smf11*, a mutant of the closest homolog of Smf12, to various cell wall-perturbing agents, including caspofungin, which inhibits glucan synthesis, and calcofluor white (CFW), which binds and perturbs the chitin layers. *smf12* together with *smf11* and their corresponding double mutant *smf12 smf11* exhibited hypersensitivity to CFW under Mn limitation as compared to their parental WT strain (**Figure 1D**). Supplementing the growth medium with Mn alleviated the CFW defect in all tested mutants, suggesting that the cell wall perturbation is a consequence of impaired Mn homeostasis. Intriguingly, in response to caspofungin, *smfs* mutants showed a moderate growth defect exclusively under Mn sufficiency while Mn depletion rescued this phenotype (**Figure 1D**). Together, this underlined that Mn homeostasis mediated by Smf12 and Smf11 is required for cell wall integrity in *C. albicans*.

### Inactivation of *SMF12* alters cell wall ultrastructure and increases **β**-glucan and chitin exposures

The regulation of the exposure of cell wall polysaccharides, a process known as masking ^28^, is crucial for the outcome of an infection, as it enables fungal pathogens to evade host immune surveillance. To assess whether the observed cell wall perturbation in *smf12* and *smf11* affect cell wall masking, we quantified by flow cytometry the exposure of β-glucan, chitin, and mannans under both Mn deprivation and sufficiency conditions. While the exposure of the three polysaccharides in WT cells was not affected by Mn availability, *smf12* and *smf11* and the corresponding double mutant exhibited a drastic increase of both β-glucan and chitin exposures when Mn was depleted (**Figure 2A-B**). The unmasking of β-glucan and chitin in these mutants was reverted by Mn supplementation suggesting that Mn homeostasis is important for the cell wall masking in *C. albicans*.

**Figure 2.**
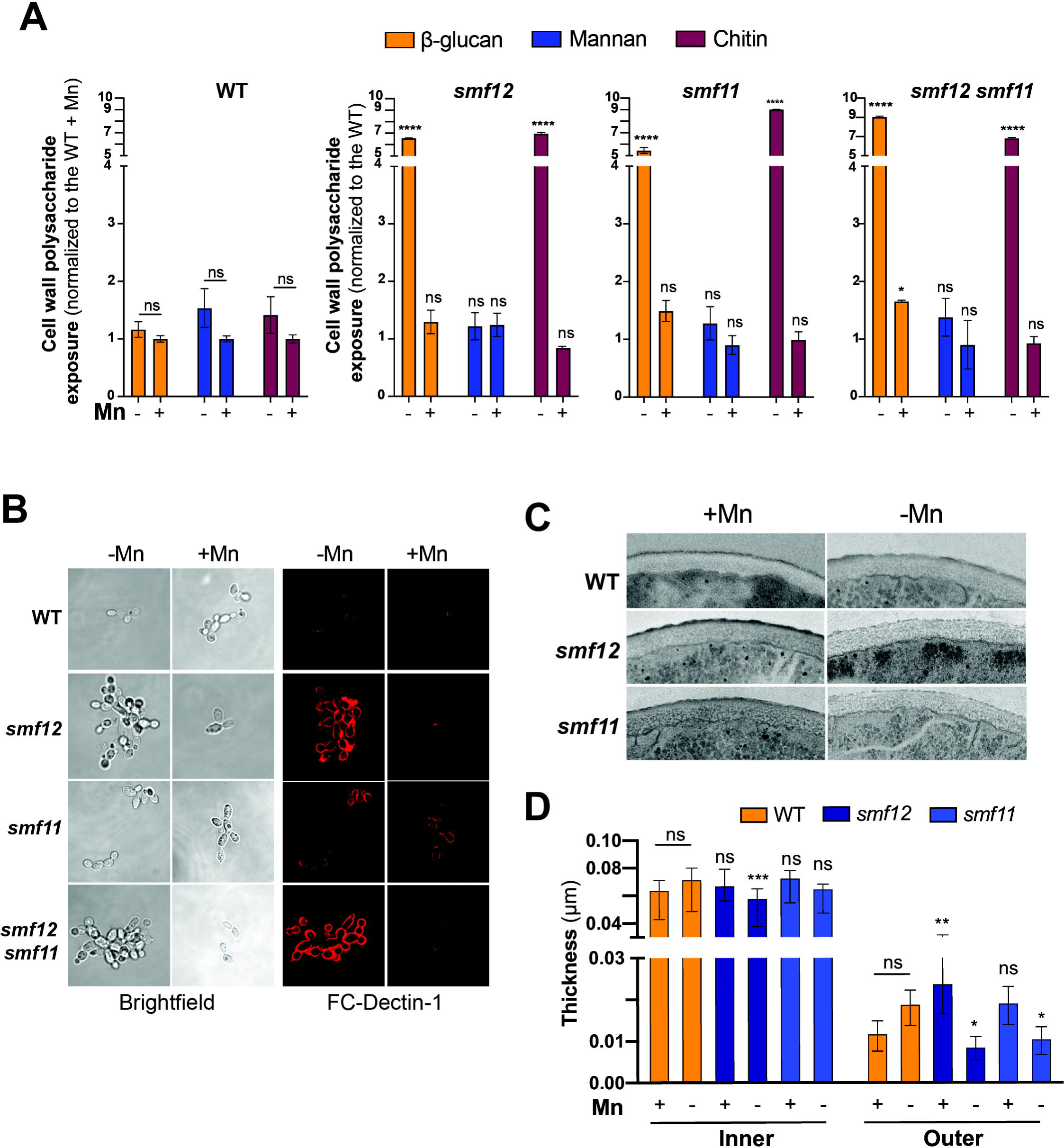
Altered cell wall ultrastructure and increased β-glucan and chitin exposures in *smf* mutants. (**A**) Quantification of β-glucan, mannan and chitin exposure by flow cytometry in WT and *smf* mutants growing under either Mn scarcity or sufficiency. Cells were collected and incubated with FC-Dectin-1, WGA-Alexa-488 and Concanavalin A-Alexa-488 to assess the β-1,3-glucan, chitin and cell wall mannan at the cell periphery, respectively. (**B**) Representative confocal microscopy image of stained of β-glucan exposure in *smf* mutants and WT strains growing in the presence and the absence of Mn. (**C-D**) Effects of genetic inactivation of *SMF12* and *SMF11* on *C. albicans* cell wall architecture. TEM was used to analyse the effect of Mn availability on cell wall ultrastructure in WT and *smf* mutants (*smf12* and *smf11*). Representative images are shown for each strain and condition(**C**). (**D**) Quantification of the thickness of the inner cell wall layer and mannoprotein fibril length form micrographs. Data represent the mean ± SEM of 100 measurements from 20 cells per condition.

The impact of *smf* mutations on the ultrastructure of the cell wall was also investigated using transmission electron microscopy. The thickness of the inner cell wall, which contains both glucan and chitin, as well as the outer cell wall with its characteristic fibrillar layer of mannoproteins, was measured. While Mn availability has no significant effect on the cell wall thickness of the WT cells, *smf12* and *smf11* showed a significant reduction of the mannan fibrillar outer layer specifically under Mn starvation which might explain the unmasking of β-glucan and chitin phenotype (**Figure 2C-D**). This supports the previous finding highlighting the crucial role of Mn homeostasis in protein mannosylation and glycosylation in *C. albicans* ^11,13^. *smf12* mutation led also to a weak but significant decrease in the inner layer thickness under Mn limitation (**Figure 2C-D**). Together, these findings support that Mn uptake deficiency led to cell wall structure perturbation and unmasking of both chitin and β-glucan.

### Cell wall unmasking in *smf12* and *smf11* under Mn scarcity affects host-pathogen interaction

Different studies have highlighted the impact of glucan and chitin exposure on the phagocytosis rate of immune cells. In general, unmasking either glucan or chitin in certain mutants or in response to specific environmental cues has been shown to enhance immune recognition and increase phagocytosis by macrophages and neutrophils ^28^. Based on this, we investigated whether the increased exposure of glucan and chitin on the surface of *smf12* and *smf11* cells led to enhanced phagocytosis by macrophages. J774A.1 murine macrophages were exposed to either WT or *smfs* mutant cells grown in an Mn-depleted or repleted medium. *C. albicans* WT strain was phagocytosed at the same rate regardless of Mn availability in the growth medium (**Figure 3A**). Unexpectedly, a significant decrease of phagocytosis rate was perceived when *smf12*, *smf11* or *smf11smf12* mutants were grown under Mn limitation (**Figure 3A**). This phagocytosis pattern was reversed when Mn was supplemented to a level comparable to that of the WT strain. This suggests that the unmasking phenotype of *smf* mutants under Mn starvation is linked to a reduced recognition or internalization by macrophages, which contrasts with previous findings ^28^.

**Figure 3.**
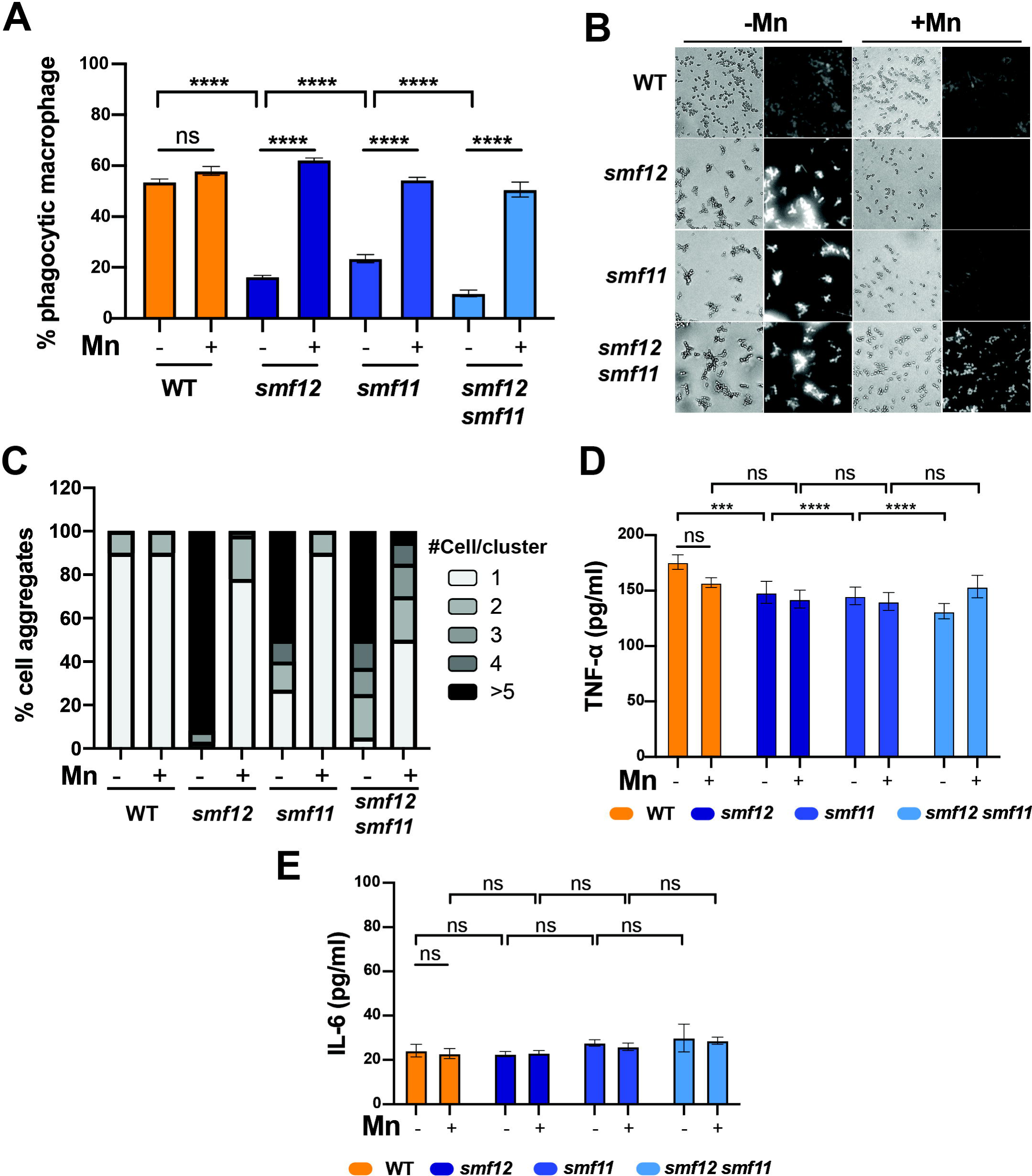
Phagocytosis of *smf* mutants by murine macrophages and the subsequent inflammatory response. (**A**) *smf* mutants and WT cells pre-grown in an Mn-depleted or-repleted medium were co-incubated with J774A.1 macrophages at an MOI of 5:1 for one hour, and the rate of phagocytosis determined by flow cytometry. For each strain and each growth condition, data represent the percentage of phagocytosis as compared to the WT in SC+Mn. (**B-C**) Altered *smf* mutants’ morphology is reverted by Mn supplementation. (**B**) Representative images of WT and *smf* mutants’ morphologies showing cells stained with CFW. (**C**) Distributions of the number of yeast cells per cluster for each strain under Mn limitation or repeltion. (**D-E**) Inflammatory responses in macrophage incubated with *smf* mutants. ELISA results are shown for TNF-α and IL-6 in supernatants of co-culture media.

The reduced phagocytosis rate in *smf* mutants under Mn starvation may be linked to their morphological defects. Most of these cells grew as larger multicellular aggregates, which could enhance resistance to phagocytosis, as previously reported for different mutants with similar morphological defect ^13,29–33^. Indeed, Mn depletion induced a clumping morphology in *smf* mutants, likely resulting from a cell separation defect, which was rescued by Mn supplementation (**Figure 3B-C**). Alternatively, the decreased phagocytosis could result from the overexposure of chitin in *smf* mutants, potentially blocking Dectin-1. Previous studies have shown that chitin interferes with Dectin-1 receptor activity, inhibiting phagocytosis and leading to reduced cytokine production and macrophage engulfment ^34–36^. Consistent with this, the overexposure of either glucan or chitin on the surface of *smf* mutants under Mn limitation did not significantly affect the immune response, as assessed by the levels of tumor necrosis factor alpha (TNF-α) and IL-6, secreted in media, compared to Mn-repletion conditions (**Figure 3D-E**). Taken together, these findings suggest that although *smf* mutants exhibit increased exposure of glucan and chitin, their reduced susceptibility to phagocytosis is likely due to their altered morphology and/or the masking of Dectin-1.

### *smf12* unmasking is not mediated by the calcineurin pathway

Gene Set Enrichment Analysis (GSEA) was used to identify similarities between the *smf12* transcriptional profile and those of potential signaling pathways that could mediate the observed unmasking phenotype. We found that *smf12* profile significantly correlates with different transcriptomics datasets of mutants of the calcineurin pathway, including *cna1* mutant of the calcineurin catalytic subunit and its major transcription factor effector, *crz1* (**Figure 4A-B and Supplementary Table S3**). A total of 36 transcripts that are activated by calcium and regulated by both calcineurin and Crz1 (60 in total) ^37^ were upregulated in *smf12* (**Figure 4C**). This set of common genes were enriched mainly in biological processes linked to cell wall remodeling, calcineurin signaling (*CRZ1*, *CCH1*, *RTA2*) stress responses and transport (**Figure 4D**). Again, supplementing the growth medium with Mn attenuated the calcineurin activation signature of *smf12* mutant suggesting a causality between Mn uptake defect and calcineurin signaling activation. Under Mn limitation, *smf12*, as well as *smf11* and *smf11smf12* mutants, exhibited elevated calcium levels compared to the WT cells, corroborating the activation of the calcineurin transcriptional output in *smf12* (**Figure 4E**).

**Figure 4.**
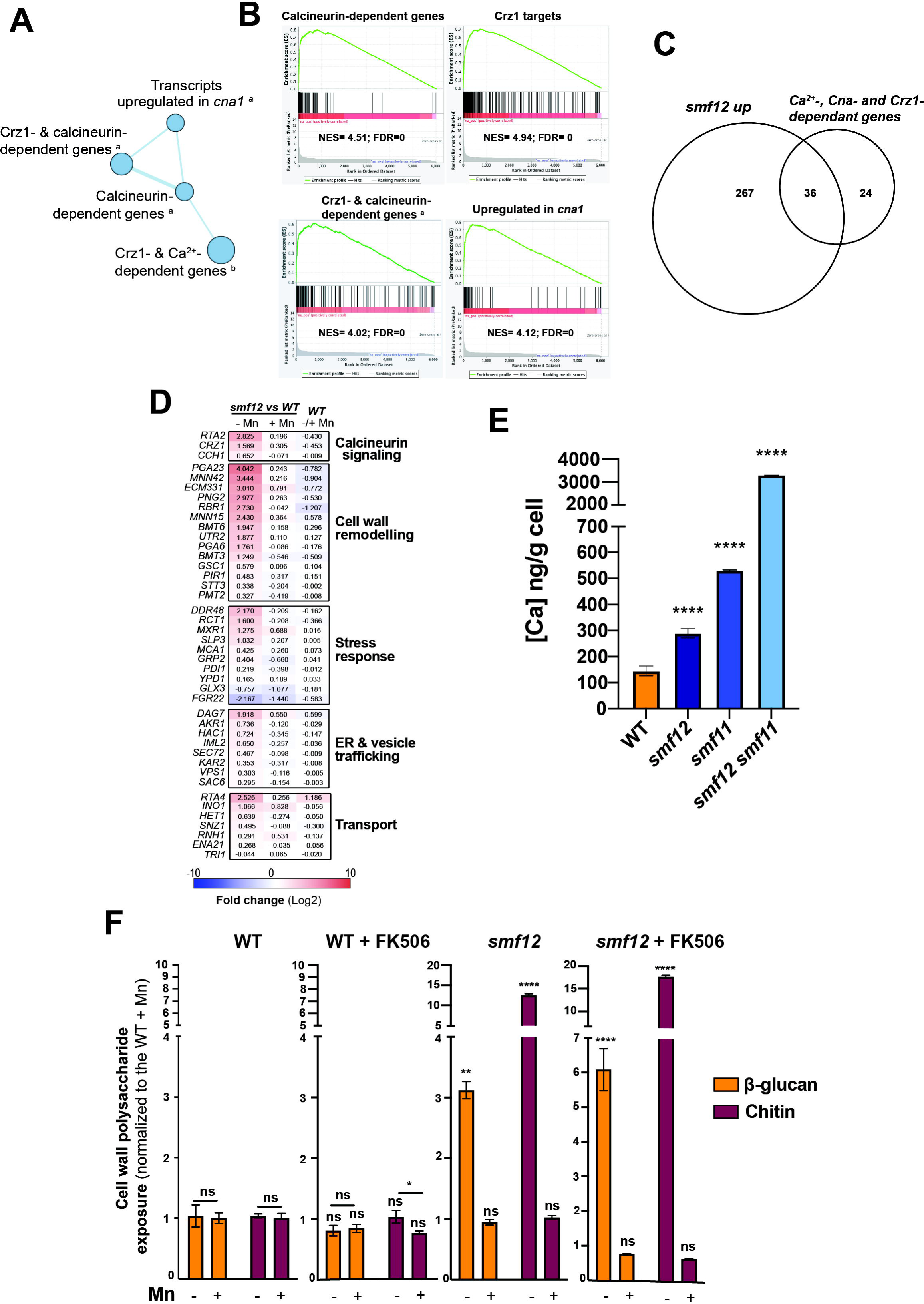
Activation of the calcineurin pathway in *smf12*. (**A**) GSEA analysis indicates a transcriptional signature reflecting calcineurin pathway activation in *smf12* mutant. The diameter of the circle reflects the number of modulated transcripts in each dataset. Images were generated using Cytoscape ^58^ with the Enrichment Map plug-in. (**B**) GSEA graphs of significant correlations between *smf12* transcriptome and mutants of the calcineurin pathway (*cna1* and *crz1*). (**C-D**) Overlap of significantly upregulated transcripts in *smf12* mutants with the set of genes defining the calcineurin regulon in *C. albicans* ^37^. (**E**) *smf* mutants exhibit high intracellular calcium (Ca) content. Ca levels were determined by ICP-MS in exponentially grown WT (SN148-CIp20) and *smf* strains in SC-Mn medium at 30°C. (**F**) *smf12* unmasking of β-glucan and chitin is not mediated by the calcineurin pathway. Quantification of β-glucan, mannan and chitin exposure by flow cytometry in WT and *smf12* growing under either Mn scarcity or sufficiency in the presence or the absence of the calcineurin inhibitor FK506.

Former study underlined the contribution of the calcineurin signaling to the unmasking phenotype of *C. albicans* ^38,39^. As our RNA-seq data reflect an activation of this signaling pathway, we hypothesized that *smf12* unmasking is mediated by the calcineurin pathway. To test this, we assessed both glucan and chitin exposures of *smf12* mutant in the presence of the calcineurin inhibitor FK506. Under Mn limitation, as described above, *smf12* exhibited an unmasking of both β-glucan and chitin (**Figure 4F**). However, inhibition of the calcineurin signaling in this mutant increased further the exposition of both β-glucan and chitin as compared to the control (**Figure 4F**). Mn repletion reverted the unmasking of *smf12* in the presence or the absence of the calcineurin inhibitor (**Figure 4F**). Thus, activation of the calcineurin signaling under Mn starvation is unlikely modulating the unmasking phenotype of *smf12*.

### *Smf* mutants exhibit a decreased Mn-dependant glucanase activity

Glucanases such as the endo-glucanase Eng1 and the exo-glucanase Xog1 were shown to mediate shaving of exposed β-glucan and contribute to the hiding of fungal pathogen from the immune system recognition ^40–42^. As our RNA-seq showed a decreased transcript level of the endo-glucanase Eng1, we hypothesized that overexposure of β-glucan on the surface of *smf12* might be a consequence of a decreased shaving capacity of this epitope. We measured β-glucanase activity of cell pellet of *smf* mutants using laminarin as a substrate. All *smf* mutants exhibited a significant decrease in glucanase activity under Mn limitation as compared to the WT strain and the Mn-repletion conditions, suggesting a reduction of the β-glucan “shaving” capacity of *smf* mutants (**Figure 5A**).

**Figure 5.**
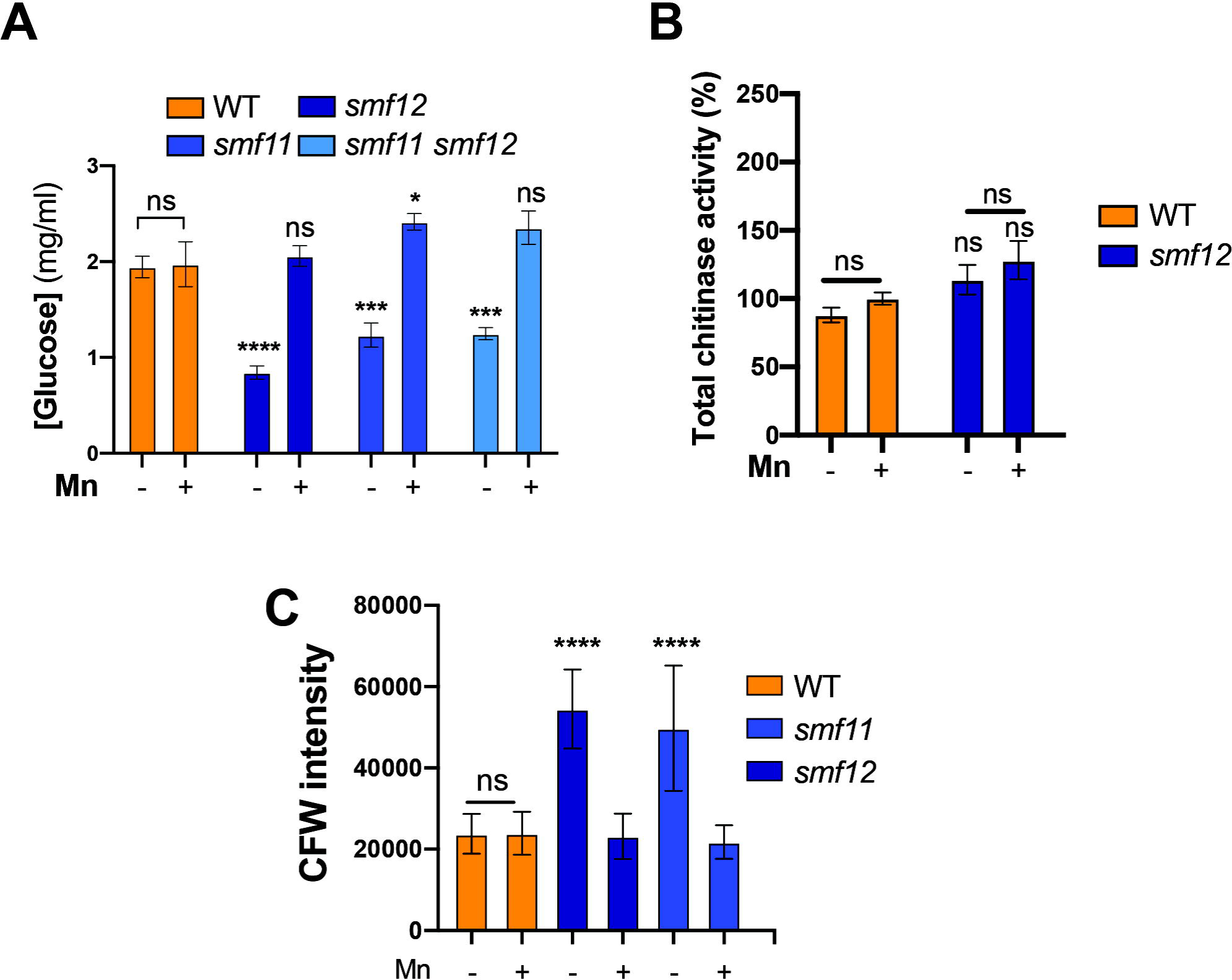
*Smf* mutants exhibit a decreased glucanase activity and a high chitin content. (**A**) Exo-glucanase activity of WT and *smf* (*smf12*, *smf11* and *smf12 smf11*) cells was determined in each strain under either Mn limitation or repletion. (**B**) Chitinase activity of WT and *smf12* in response to Mn availability. (**C**) Total chitin assessment in WT, *smf12* and *smf11* strains. Total chitin was quantified using microscopy and CFW staining. For each condition, fluorescence intensities of 100 cells were measured using ImageJ.

### Chitin synthesis, but not processing, is altered in *smf12*

Previous work have shown that chitin unmasking in response to low pH resulted from a reduced expression of the cell wall chitinase Cht2 leading to decreased processing of chitin fibrils ^43^. As our RNA-seq data showed a downregulation of *CHT2* and also *CHT3,* we hypothesized that a similar mechanism causes de-cloaking of chitin at the cell periphery of *smf12* under Mn limitation. When comparing chitinase activity of the WT and *smf12* strains under either Mn scarcity or repletion no significant difference was noticed (**Figure 5B**). Furthermore, other studies highlighted the role of increased chitin biosynthesis and deposition at the unmasked foci of the cell wall in altering its architecture, which consequently leads to the overexposure of glucan ^39,44^. Indeed, *smf12* exhibited elevated transcript levels of the chitin synthase *CHS2* and an increased cellular chitin content, particularly under Mn limitation. (**Figure 5C**). Collectively, these findings indicate that the chitin unmasking observed in *smf12* is likely due to enhanced chitin biosynthesis and deposition at the cell wall, rather than a defect in chitin cleavage.

## Discussion

Recent studies have highlighted the critical role of Mn homeostasis in regulating various biological processes that influence the fitness and virulence of *C. albicans* ^10–12^. Mn has been shown to play an important role in *C. albicans*’ ability to cope with both endoplasmic reticulum and plasma membrane stress, and it is essential for the functionality of superoxide dismutase enzymes. In this study, we uncovered a novel function of Mn in maintaining cell wall integrity and modulating the exposure of fungal antigenic determinants, further emphasizing the central role of this metal in supporting the opportunistic nature of this yeast.

The sensitivity of *smf* mutants to CFW underscores the critical role of Mn metabolism in maintaining the integrity of the *C. albicans* cell wall. This defect is reflected in the transcriptome of *smf12*, with differential regulation of many genes involved in cell wall biogenesis, as well as transcriptional signatures indicative of the activation of cell wall integrity (CWI) pathways, including calcineurin and the MAPK Cek1. The cell wall defect observed in *smf12* could be primarily attributed to alterations in the abundance of chitin on the cell surface. Notably, high chitin content in *C. albicans* and other *Candida* species has previously been linked to hypersensitivity to CFW as shown here for *smf* mutants ^45^. Conversely, increased chitin content has been associated with enhanced tolerance to caspofungin ^45–47^. This might explain why *smf* mutants exhibit sensitivity to this echinocandin only when Mn is available, whereas under Mn-limited conditions, the elevated chitin content may help mitigate their sensitivity to caspofungin. Furthermore, the mannosylation defect observed in *smf* mutants, as previously reported ^10,11^, together with the reduced thickness of the internal cell wall layer reported here, could lead to increased chitin levels, similar to what was seen in mannosyltransferase mutants in *C. albicans*^48^_._

In this study, we demonstrate that impairing *C. albicans* cells’ ability to uptake Mn results in the unmasking of both chitin and β-glucan. This phenotype is likely the result of a combination of various alterations in the *smf* mutants. First, ultrastructural analysis of the cell wall showed a reduction of the mannan fibrillar outer layer which might primarily contribute to the unhiding of β-glucan from the cell surface as previously observed in *C. albicans* mannosylation- and mannoprotein-deficient mutants ^49–51^. Second, increased chitin content and deposition at the cell surface may drive β-glucan unmasking in *smf12*. Indeed, β-glucan unmasking in *C. albicans*, whether triggered by antifungal drugs, environmental or host signals, is typically accompanied by increased chitin deposition at the cell periphery ^28^. Thus, abnormal deposition of chitin layer at the cell wall of *smf12* might push β-glucan to emerge at the cell periphery. Third, while chitin processing by chitinases was unaffected in *smf12* and *smf11*, glucanase activity was significantly impaired, leading to a reduced capacity for β-glucan “shaving” in these mutants. Notably, the transcript levels of the endo-glucanase Eng1 were reduced in *smf12*, suggesting that the unmasking of β-glucan could result from improper modulation of these enzymes and an inability to cleave exposed β-glucan from the cell surface ^40^. Overall, it is likely that one or more of these mechanisms, or a combination thereof, explain the unmasking of β-glucan and chitin in *smf* mutants. Overall, it may be possible that any one of the above mechanisms, and more than likely many of them, explain the unmasking of β-glucan and chitin in *smf* mutants.

In *C. albicans*, β-glucan unmasking is regulated by multiple signaling mechanisms including the calcineurin and the Cek1 MAPK pathways ^28,38^. In the current study, although calcineurin was activated in *smf12*, as evidenced by the activation of its *bona fide* regulon, inhibition of this signaling pathway failed to restore β-glucan and chitin exposure to wild-type levels. This suggests that the unmasking phenotype in *smf12* is independent of the calcineurin pathway. Our RNA-seq data revealed an upregulation of the MAPK Cek1, indicating a potential role for this signaling pathway in mediating the unmasking phenotype observed in *smf12*. However, GSEA analysis did not reveal a significant similarity between the *smf12* profile and mutants of the Cek1 pathway (e.g. mutant of the transcription factor effector, Cph1 and *STE11*^ΔN467^ hyperactive mutant). Therefore, further investigation is required to determine whether Cek1 or another, yet unidentified, pathway is mediating the unmasking phenotype under Mn deficiency. Additionally, the cause of calcineurin hyperactivation in *smf12* remains to be elucidated. In many fungi, the calcineurin pathway is a well-known regulator of both cell wall integrity and UPR ^52–54^. Accordingly, one possible explanation is that the calcineurin pathway could be activated as a secondary response to signal cell wall alterations and/or ER stress, given that the UPR is constitutively activated in *smf12* ^55,11^. Furthermore, calcineurin activation might prevent the formation of non-viable supra-high chitin *smf12* cells by modulating chitin biosynthesis as previously reported ^56^.

An unexpected finding of the current study is that the unmasking of either chitin or glucan did not lead to an enhanced immune response or increased phagocytosis, as initially anticipated. The reduced susceptibility of *smf* mutants to phagocytosis is likely due to their altered morphology, as they formed aggregates, presumably resulting from a defect in chitin processing at the septa. Similar results have been observed in *C. albicans* phosphomannan-deficient mutants and other mutants of glycosyltransferases, which also exhibit an aggregate phenotype akin to that of *smf12* and show resistance to phagocytosis ^30,32^. Recent studies on the multi-drug-resistant yeast *C. auris* have demonstrated that evolved aggregative cells, or cells where aggregation is induced by specific drugs or growth conditions, are more resistant to phagocytosis by macrophages than their yeast-form counterparts ^29,57^. Future research will be essential to further clarify the role of Mn homeostasis in the development of this multicellular morphology and to investigate whether Mn-starved niches within the human host may contribute to the immunity-recalcitrant phenotype observed in *C. auris*. The decreased uptake of *smf* mutants by macrophages may be a consequence of Dectin-1 masking, likely due to the high chitin content on their cell surface. Interestingly, this fungal survival strategy has been recently described in *C. albicans*, where increased chitin on the cell surface promotes arginine degradation by host arginase-1, thereby preventing the utilization of this amino acid by nitric oxide synthase to produce the antifungal molecule nitric oxide ^34^.

## Disclosure statement

No potential conflict of interest was reported by the author(s).

## Funding

This work was supported by grants from the Canadian Institutes of Health Research (Project grant), the Canada Foundation for Innovation and the Montreal Heart Institute foundation Startup fund to AS. AS is supported by a Fonds de Recherche du Québec-Santé Senior Salary award. MH is supported by a Natural Sciences and Engineering Research Council of Canada (NSERC) CREATE PhD scholarship (EvoFunPath program).

## Data availability statement

All RNA-seq data are available at the GEO database (https://www.ncbi.nlm.nih.gov/geo/) under the accession number GSE283948. The data that support the findings of this study are openly available in https://doi.org/10.6084/m9.figshare.28114337.v1.

## Author contributions statement

MH and AS were involved in the conception, design, analysis, interpretation of the data; They also drafted and revised the final version of the manuscript. MH, MK, GTJ, LV and ATV carried out the experiments and analysed the results. ER, JCT and ATV revised critically the manuscript for intellectual content. All authors agree to be accountable for all aspects of the work.

## Supplementary data

**Supplementary Table S1. Strains and primers used in this study.**

**Supplementary Table S2. *smf12* RNA-seq data.**

**Supplementary Table S3. GSEA analysis of the *smf12* transcriptome.**

## Notes

### Competing Interest Statement

The authors have declared no competing interest.

